# Targeted Whole Genome Sequencing of African Swine Fever Virus and Classical Swine Fever Virus on the MinION Portable Sequencing Platform

**DOI:** 10.1101/2025.07.16.665162

**Authors:** Chester D. McDowell, Taeyong Kwon, Patricia Assato, Emily Mantlo, Jessie D. Trujillo, Natasha N. Gaudreault, Leonardo C. Caserta, Igor Morozov, Jayme A. Souza-Neto, Roman M. Pogranichniy, Diego G. Diel, Juergen A. Richt

## Abstract

African swine fever virus (ASFV) and classical swine fever virus (CSFV) are important transboundary animal diseases (TADs) affecting swine. ASFV is a large DNA virus with a genome size of 170-190 kilobases (kB) belonging to the family Asfarviridae, genus Asfivirus. CSFV is a single-stranded RNA virus with genome size of approximately 12 kB belonging to the family Flaviviridae, genus Pestivirus. Outbreaks involving either one of these viruses result in similar disease syndromes and significant economic impacts from: (i) high morbidity and mortality events; (ii) control measures which include culling and quarantine; and (iii) export restrictions of swine and pork products. Current detection methods during an outbreak provide minimal genetic information on the circulating virus strains/genotypes that are important for tracing and vaccine considerations. The increasing availability and reduced cost of next-generation sequencing (NGS), allows for the establishment of vital NGS protocols for the rapid identification and complete genetic characterization of outbreak strains during an investigation. NGS data provides a better understanding of viral spread and evolution facilitating the development of novel and effective control measures. In this study, panels of primers spanning the genomes of ASFV and CSFV were independently developed to generate approximately 10kB and 6kB amplicons, respectively. The primer panels consisted of 19 primer pairs for ASFV and 2 primer pairs for CSFV providing whole genome amplification of each pathogen. These primer pools were further optimized for batch pooling and thermocycling conditions, resulting in a total of 5 primer pools/reactions used for ASFV and 2 primer pairs/reactions for CSFV. The ASFV primer panel was tested on viral DNA extracted from blood collected from pigs experimentally infected with ASFV genotype I and genotype II viruses. The CSFV primer panel was tested on 11 different strains of CSFV representing the 3 known CSFV genotypes, and 21 clinical samples collected from pigs experimentally infected with 2 different genotype 1 viruses. ASFV and CSFV amplicons from optimized PCR reactions were subsequently sequenced on the Oxford Nanopore MinION platform. The targeted protocols for these viruses resulted in an average coverage greater than 1000X for ASFV with 99% of the genome covered, and 10,000X-20,000X for CSFV with 97% to 99% of the genomes covered. The ASFV targeted whole genome sequencing protocol has been optimized for genotype II ASF viruses that have been responsible for the more recent outbreaks outside of Africa. The CSFV targeted whole genome sequencing protocol has universal applications for the detection of all CSFV genotypes. Protocols developed and evaluated here will be essential complementary tools for early pathogen detection and differentiation as well as genetic characterization of these high consequence swine viruses, globally and within the United States, should an outbreak occur.

## 1. Introduction

Transboundary animal diseases (TADs) are of great concern to the global agricultural systems. TADs can easily traverse national and international borders resulting in devasting economic impacts due to high morbidity and mortality events, costly control strategies, and imposed trade restrictions. This study focuses on two TADs that have significant impacts on the global pork industry. One of these TADs currently in the spotlight is African swine fever virus (ASFV) which is the causative agent of African swine fever, a contagious viral disease of domestic swine and wild boar (Sus Scrofa) that can cause high mortality in affected swine populations. ASFV belongs to the Asfarviridae family, genus Asfivirus [1,2]. The virus particle of ASFV encapsulates a 170-192 kB double stranded DNA genome encoding more than 150 genes [3–6]. Historically, there are 24 genotypes of ASFV with varying virulence between and within genotypes; recent retrospective studies of ASFV sequences suggest that there are only 6 to 7 ASFV genotypes including the genotype I/II virus recombinant virus isolated in China [7,8]. Since the 2007 outbreak in Georgia, ASFV’s global presence has expanded to central and eastern Europe, Asia, and the Caribbean [9]. Another important TAD is classical swine fever virus (CSFV), the causative agent of classical swine fever or historically termed “hog cholera” [10]. CSFV belongs to the Flaviviridae family, genus Pestivirus; the enveloped virus particle contains a non-segmented single-stranded positive sense RNA genome of approximately 12 kB en-coding 13 viral proteins [11,12]. Currently there are three known genotypes with over 13 sub-genotypes [13]. CSFV was first described in the United States and has since been eradicated from the U.S. swine population. However, CSFV remains endemic in many countries around the world. The risk of introduction and reintroduction into currently CSFV-free countries remains high. Until these two TADs are controlled and eventually eradicated, they will continue to threaten the global pork industry. Currently, rapid detection and implementation of strict mitigations strategies is the most effective way of impeding global spread.

Currently, the World Organization for Animal Health (WOAH) recommends the use of qPCR or RT-qPCR for the detection of ASFV and CSFV, respectively [14,15]. These methods offer high specificity and sensitivity with shorter turnaround times compared to traditional diagnostic techniques such as antigen ELISA and virus isolation. However, these techniques are unable to provide genotypic information of circulating viral strains in an outbreak. Traditionally, this information is obtained by sequencing specific genes or regions that are conserved within genotypes. In the case of ASFV, genotyping is based on the sequencing of the C-terminal variable region of the B646L gene that encodes the major capsid protein, p72 (478 bp) [16,17]. ASFV strains are further characterized by sequencing of the envelope protein (p54), the central hypervariable region (CVR) of B602L (viral chaperone), and the CD2v (EP402R) and C-type lectin genes (EP153R) [18,19]. Genotyping CSFV is achieved by sequencing the 5’-NTRs, the partial or full E2 gene, the partial NS5B gene, or the whole viral genome [15,20]. Due to the size of the ASFV genome and the cost of sequencing in previous years, partial sequencing of these two agents was sufficient for genetically characterizing circulating or outbreak strains. However, partial sequencing provided only limited genetic information, has the potential for erroneous characterization of viral strains and limits researchers’ ability to fully understand the viral evolution of these pathogens [7]. Recent studies by Forth et al. investigating the genetic landscape of ASFV strains circulating in Germany found that there are multiple variants that have di-verged from the original outbreak strain [21].

The rapid advances in sequencing technologies have greatly reduced the cost and time needed to obtain whole genome sequences of these important pathogens and other viral agents. The development of the Oxford Nanopore MinION, a portable sequencing platform, has drastically changed the landscape of sequencing. The MinION platform us-es a pore-based sequencing technology that has eliminated read length restrictions observed with previous sequencing technologies. This advancement has allowed for the development of targeted long read amplicon-based sequencing protocols. The MinION plat-form is relatively inexpensive compared to the bench top sequencing platforms, increasing the availability of sequencing technology [22].

In this study we aimed to develop targeted whole genome sequencing protocols using the MinION sequencing platform for the complete genetic characterization of ASFV and CSFV. The protocols are designed to rapidly obtain whole genome sequences of these two viruses from clinical samples within 36 to 48 hours. These protocols will aid in increasing the availability of whole genome sequences of ASFV and CSFV and will support efforts to provide additional genetic information to better understand viral evolution and aid in the development of novel diagnostics and effective vaccines.

## 2. Materials and Methods

### 2.1 Samples Ethics Statement

All experiments involving African swine fever virus (ASFV) and classical swine fever virus (CSFV) were conducted in the BSL-3 laboratory at the Biosecurity Research Institute (BRI) at Kansas State University (KSU) in Manhattan, KS. All animal studies and experiments were approved and performed under the KSU Institutional Biosafety Committee (IBC, Protocol #s:1651, 1668 and 1787) and the Institutional Animal Care and Use Committee (IACUC, Protocol #s: 4265 and 4363) in compliance with the Animal Welfare Act. The ASFV and CSFV clinical samples used in this study were acquired from experimentally infected pigs at KSU-BRI and cell culture derived virus stocks (Tables 3, 4, and 5).

### 2.2 Nucleic acid Extraction and Quantitative Real-time PCR

#### 2.2.1 African Swine Fever Virus DNA Extraction and Quantitation

ASFV specific DNA was detected and quantified in EDTA blood and spleens from experimentally infected pigs with ASFV strains Armenia/2007, Mongolia/2019 [23], or E70 by quantitative real-time PCR (qPCR) assay as previously described [23,24]. ASFV DNA was extracted using two different automated extraction systems, one for EDTA blood and one for spleen. For EDTA blood, total DNA was extracted using the MagMax Pathogen RNA/DNA kit (ThermoFisher Scientific, Waltham, MA, USA) using the Kingfisher™ Duo Prime purification system, an automated magnetic bead extraction system, following the manufacturer’s protocol with minor modifications. Briefly, EDTA blood was mixed 1:1 with Buffer AL (Qiagen; Carlsbad, CA, USA) and heated at 70 °C for 10 min. Subsequently, 200 µL was mixed with 20 µL of bead mix consisting of an equal volume of nucleic acid binding beads and lysis enhancer. The mixture was gently mixed by pipetting and incubated for 2 min at room temperature. Phosphate-buffered saline (PBS, 100 µL) was added to the tube and the entire volume was added to the extraction plate well containing 400 µL lysis buffer, binding buffer and isopropyl alcohol. DNA bound beads were washed twice with 300 µL of wash solution 1, once with 450 µL wash solution 2, and once more with 450 µL of 200 proof molecular grade ethanol. The DNA bound beads were dried for 5 min prior to being eluted in 100 µL elution buffer. Negative (molecular grade water/PBS) and positive (ASFV p72 plasmid) controls were included with each extraction.

The spleen samples were processed into 20% w/v tissue homogenates in Buffer ATL (Qiagen; Carlsbad, CA, USA) with a subsequent 30-minute Proteinase K (80µg, Qiagen; Carlsbad, CA, USA) digestion at 56°C prior to nucleic acid extraction. Total DNA was extracted from tissue homogenates using the GeneReach Taco Mini Automatic Nucleic Acid Extraction System (GeneReach Biotechnology Corp., Taichung, Tiawan). Briefly, tissue homogenates were mixed at 1:1 ratio with Buffer AL. Then, 200 µl of tissue homogenate and Buffer AL mixture were added to extraction plate wells containing 50µl of magnetic, 500µl lysis buffer, 100µl phosphate buffered saline (PBS), followed by the addition of 200µl isopropyl alcohol to each well. DNA bound beads were washed twice with 750µl of wash buffer A, once with 750µl wash buffer B, and once more with 750µl of 200 proof molecular grade ethanol. The DNA bound beads were dried for 5 minutes prior to being eluted in 100µl elution buffer. Negative (molecular grade water/PBS) and positive (ASFV p72 plasmid) controls were included with each extraction.

The ASFV DNA extracted from EDTA blood and spleens was detected and quantified using the primers and probe for the detection of the p72 ASFV gene as previously de-scribed by Zsak et al. [25], using a validated protocol performed with the PerfeCTa® Fast-Mix® II (Quanta Biosciences, Gaithersburg, MD, USA) on the CFX96 Touch™ Real-Time PCR Detection System (Bio-Rad, Hercules, CA, USA) as previously described [26]. Duplicate qPCR reactions were performed. Each PCR run included negative and positive controls which consisted of nuclease-free molecular grade water and ASFV positive amplification control (p72 plasmid), respectively.

#### 2.2.2 Classical Swine Fever Virus RNA Extraction and Quantitation

Specific CSFV RNA was detected and quantified from the serum, tonsils, and a mandibular lymph node of pigs experimentally infected with either Brescia or Bavaro strains of CSFV and from cell culture supernatant of 11 CSFV viral stocks by quantitative real time reverse transcriptase PCR (RT-qPCR) described by Eberling et. al [27]. Prior to nucleic acid extraction, tissue homogenates were prepared from the tonsils and a mandibular lymph node using the same method as for ASFV spleens described above. Viral RNA was extracted from cell culture supernatant, serum, and tissue homogenates using the Gene-Reach Taco Mini Automatic Nucleic Acid Extraction System (GeneReach Biotechnology Corp., Taichung, Tiawan) as described above for the ASFV spleen samples except with Buffer AL substituted by Buffer RLT (Qiagen; Carlsbad, CA, USA). Negative (molecular grade water/PBS) and CSFV positive extraction controls (CSF phagemid provided by USDA-APHIS FADDL) controls were included with each extraction.

The CSFV RNA extracted from cell supernatant, serum, mandibular lymph node, and tonsils was detected and quantified using the primers and probe for detection of CSFV de-scribed by Eberling et.al. using a validated protocol performed with qScript XLT 1-step RT-qPCR ToughMix (Quanta Biosciences, Gaithersburg, MD, USA) on the CFX96 Touch™ Real-Time PCR Detection System (Bio-Rad, Hercules, CA, USA). Briefly, the final reaction mixture consisted of 2.5 µl of viral RNA combined with 17.5 µl master mix including 0.2 µM forward primer, 0.4 µM reverse primer, and 0.2 µM probe. The thermocycling conditions were as follows: 50ºC for 20 min, 95ºC for 5 min, followed by 45 cycles at 95ºC for 10 sec and 60ºC 1 min. Duplicate qPCR reactions were performed. Each PCR run included negative and positive controls which consisted of nuclease-free molecular grade water and CSFV positive amplification control (CSF phagemid supplied by USDA-APHIS FADDL), respectively.

### 2.3 Primer Design

#### 2.3.1 African Swine Fever Virus

A panel of 19 ASFV primer pairs (Table 1 and Figure 1) were designed based on the ASFV genotype II Georgia 2007 genome using Primer-BLAST (NCBI), Snapgene (GSL Bio-tech LLC, Boston, MA, USA) and CLC Genomic Workbench 23.0.1 (Qiagen; Carlsbad, CA, USA). The tiled primers were designed to generate approximately 10 kB amplicons with 150-200 base pair (bp) overlaps at the 3’ and 5’ ends. All primers were designed with similar melting temperatures allowing for the use of one set of thermocycling conditions for all 19 reactions and primer pooling applications.

**Table 1.**
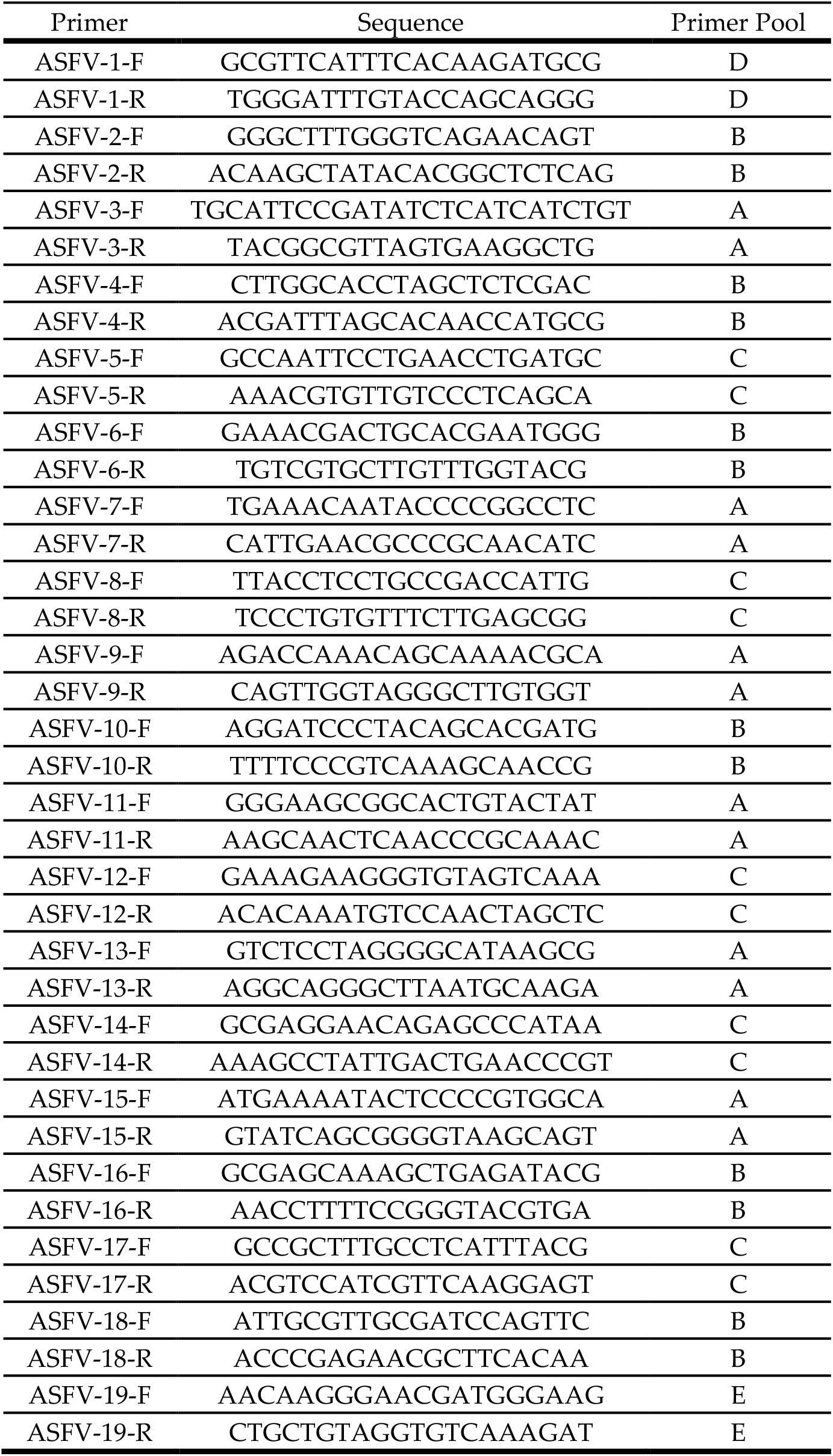
ASFV Primers and associated pools.

**Figure 1.**
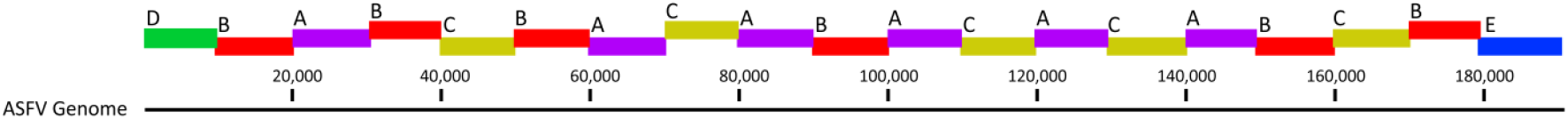
Genome map of the ASFV genome displaying the nineteen 10 kB amplicons. The colored bars represent the 5 primer pools: A: purple, B: red, C: yellow, D: green, E: blue.

#### 2.3.2 Classical Swine Fever Virus

A panel of 2 CSFV primer pairs (Table 2) were designed using CLC genomics work-bench 23.0.1 (Qiagen; Carlsbad, CA, USA) to generate approximately 6 kB fragments with a 380 bp overlap (Figure 2). The primer design module within the CLC genomic work-bench 23.0.1 (Qiagen; Carlsbad, CA, USA) was used to identify candidate primers. The candidate primer sequences were compared to an alignment of multiple CSFV genome sequences to confirm binding sites were in conserved or low diversity regions within the CSFV genome. Degenerate nucleotides were added to primer sequences in regions of diversity for amplification of different CSFV genotypes. All primers were designed to have similar melting temperatures so thermocycling conditions were consistent for the 2 primer pairs.

**Table 2.**
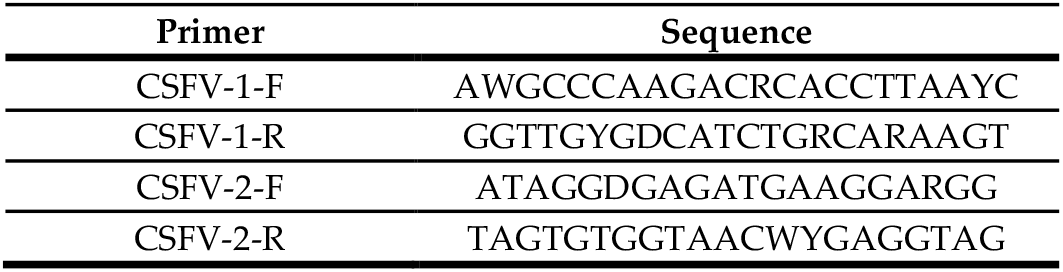
CSFV Primers.

**Figure 2.**
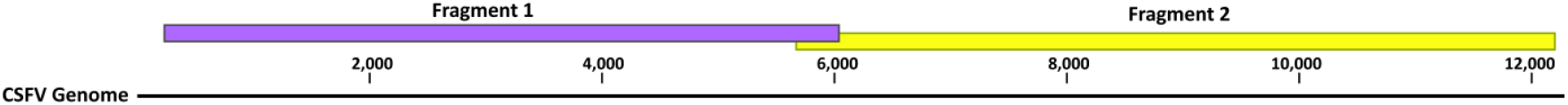
Genome map of CSFV genome displaying the two approximately 6 kB amplicons generated from the two primer pairs.

### 2.4 Whole Genome Amplification

#### 2.4.1 African Swine Fever Virus

PCR reactions for all 19 primer pairs were tested individually for specificity and optimization of thermocycling condition using ASFV viral DNA isolated from experimentally infected pig samples described above. Individual primer sets tested in 20 µl PCR re-actions consisted of 10 µl Platinum Superfi II Green PCR MasterMix (ThermoFisher Scientific, Waltham, MA, USA), 1 µl of primer pools (20 µM), 2 µl of viral DNA, and molecular grade water. The following thermocycling conditions were used: 98oC for 30 seconds, followed by 35 cycles at 98oC for 10 seconds, 60oC for 10 seconds, 68oC for 5 minutes, fol-lowed by a final extension step of 72oC for 5 minutes. Correct amplicon sizes were con-firmed by gel electrophoresis. Once optimal thermocycling conditions were achieved, 5 primer pools (pools A-E) were optimized to reduce the number of reactions needed for whole genome amplification (Figure 1); primer sequences and associated primer pools are provided in Table 1. ASFV amplicons were generated using the 5 primer pools using the following 40 µl PCR reactions consisting of 20 µl Platinum Superfi II Green PCR MasterMix (ThermoFisher Scientific, Waltham, MA, USA), 4 µl of primer pools (20 µM), 4 µl of viral DNA, and molecular grade water. The following thermocycling conditions were used: 98oC for 30 seconds, followed by 35 cycles at 98oC for 10 seconds, 60oC for 10 seconds, 68oC for 5 minutes, followed by a final extension step of 72C for 5 minutes. Amplicon pools were individually purified using Beckman Coulter AMPure XP beads (Beck-man Coulter, Indianapolis, IN, USA) following the manufacturer’s protocol. Following the PCR clean-up with AMPure beads, the DNA concentration for each amplicon pool was quantified using the Invitrogen Qubit 4 Fluorometer (ThermoFisher Scientific, Waltham, MA, USA) with the broad range dsDNA quantification kit. The five amplicon pools A-E were combined into one tube at a 1:1:1:1:1 ratio. The final pool was used for sequencing library preparation.

#### 2.4.2 Classical Swine Fever Virus

A two-step RT-PCR was used to amplify the two 6 kB amplicons. Firstly, viral cDNA was generated for each fragment individually using the LunaScript RT Master Mix kit (primer-free) (New England Biolabs, Ipswich, MA, USA) using a 20 µl reaction consisting of 4 µl of LunaScript RT Master mix, 1 µl of CSFV reverse primer (10 µM), 5 µl of viral RNA, and molecular grade water. The reaction was completed using the following thermocycling conditions: 55oC for 10 minutes followed by 95oC for 1 minute. Following cDNA synthesis 5 µl of viral cDNA was added to a 45 µl PCR master mix consisting of 25 µl of Q5 High-Fidelity 2X Master Mix (New England Biolabs, Ipswich, MA, USA), 5 µl of primer pools (10 µM), and molecular grade water. The reaction was completed using the following thermocycling conditions: 98oC for 1 minute, followed by 35 cycles at 98oC for 10 seconds, 60oC for 30 seconds, 72°C for 5 minutes, followed by a final extension step of 72oC for 5 minutes and hold at 4oC. Amplicon size was confirmed by gel electrophoresis. Amplicons were individually purified using Beckman Coulter AMPure XP beads (Beck-man Coulter, Indianapolis, IN, USA) following the manufacturer’s protocol. Following the PCR clean-up with AMPure beads, the DNA concentration for each amplicon was quantified using the Invitrogen Qubit 4 Fluorometer (ThermoFisher Scientific, Waltham, MA, USA) with the broad range dsDNA quantification kit. The two amplicons were combined into one tube at a 1:1 ratio. The final pool was used for sequencing library preparation.

### 2.5 Sequencing Library Preparation and MinION Sequencing

Sequencing libraries for both ASFV and CSFV were prepared using the Oxford Nanopore Rapid Barcoding Kit 96 (Oxford Nanopore Technologies, OX4 4DQ, UK) following the manufacturer’s protocol. For ASFV, sequencing libraries consisting of 6 barcoded samples were prepared. For CSFV, sequencing libraries consisting of up to 24 barcoded samples were prepared. Barcoded libraries were loaded onto the Oxford Nanopore Min-ION R9 flowcell (Oxford Nanopore Technologies, OX4 4DQ, UK) following manufacturer’s protocol. The MinION was run for 16 hours with a read length inclusion of greater than 1 kB. Following completion of the run, base calling and demultiplexing was completed using the Oxford Nanopore MinKNOW software (Oxford Nanopore Technologies, OX4 4DQ, UK). Parsed reads were imported into CLC genomics workbench 23.0.1 (Qiagen; Carlsbad, CA, USA). ASFV and CSFV sequencing reads were mapped to their respective reference genomes for coverage graph generation and mapping statistics. For several CSFV strains de novo assembly was completed and contigs were aligned using BLAST to determine the most similar available CSFV genome which served as the reference sequence for the analysis described above.

## 3. Results

### 3.1 ASFV AmpliSeq

The protocol was tested on 20 different ASFV samples. The samples included 19 clinical samples collected from pigs experimentally infected with ASFV genotype I E70 virus, ASFV genotype II Armenia/2007 and Mongolia/2019 viruses, as well as Armenia/2007 propagated in cell culture (see Table 3). The selected ASFV samples had Ct values for the ASFV p72 gene ranging from 14-33; samples with a Ct of less than 25 were di-luted 1:10 to prevent inhibition of the PCR reaction. Sequencing reads for genotype II samples were mapped to their respective ASFV genomes, Armenia/2007 or Mongolia/2019 (Figure 3). Sequencing reads for the ASFV genotype I E70 sample were mapped to a simi-lar genotype I virus, E75. The average coverage of read mapping was greater than 1,000X, covering 99% of the ASFV genome except for samples from animals #304, 190, and 79 (Table 3). Animal 304 had a high Ct value which may have contributed to the drop in cover-age. Furthermore, the DNA samples may have been degraded from freeze/thaws necessary to perform other assays. Overall, following the 16-hour MinION runs with 6 samples per run, on average 220,000 sequencing reads were obtained with an average read length of 2.5 kB and 72% of reads mapping to the reference genome. For the genotype II viruses ASFV Armenia 2007 and Mongolia 2019, 99% of the reference genome was covered with an average coverage of 1,764X. For ASFV genotype I E70 strain, ∼59,000 reads mapped to 85% of the genome were covered with an average coverage of 1,081X. The regions that were not amplified and sequenced ranged in size from 8,000-20,000 bp with the largest region at the 3’ end which is a region that encodes numerous multiple gene families (MGFs). These regions may not have been amplified due to sequence differences at the primer binding sites. The ASFV AmpliSeq protocol obtained the near full-length genome of genotype II viruses with 99% coverage and ∼1000X coverage for the tested samples. The ASFV AmpliSeq protocol will allow researchers to investigate the evolution of genotype II viruses in detail which have resulted in the most recent ASFV global outbreaks. Importantly, this protocol will provide sufficient genetic information for identifying genotype I/II recombinant ASFV viruses which have recently been identified.

**Table 3.** Sequencing data for ASFV samples tested with ASFV AmpliSeq.

| Animal ID | DPC | Type | Strain | Genotype | Ct | Reference Length | Mapped Reads | Total Reads | % Reads Mapped | Average Coverage | % Reference Covered | Average Read Length |
| --- | --- | --- | --- | --- | --- | --- | --- | --- | --- | --- | --- | --- |
| Arm07 Cos-1 | P3 | Cell culture | Arm07 | II | 27 | 190,584 | 51,134 | 65,081 | 78.57 | 1,017.61 | 0.99 | 3,749 |
| 63 | 7 | Blood | Arm07 | II | 16 | 190,584 | 127,541 | 158,104 | 80.67 | 2,138.01 | 0.99 | 3,157 |
| 67 | 9 | Blood | Arm07 | II | 19 | 190,584 | 149,958 | 184,043 | 81.48 | 2,323.76 | 0.99 | 2,928 |
| 87 | 7 | Blood | E70 | I | 25 | 181,187 | 59,820 | 91,435 | 97.37 | 1,081.24 | 0.85 | 3,357 |
| 190 | 7 | Blood | Arm07 | II | 15 | 190,584 | 60,571 | 80,273 | 75.46 | 864 | 0.99 | 2,681 |
| 192 | 7 | Blood | Arm07 | II | 16 | 190,584 | 85,585 | 112,479 | 76.09 | 1,226.02 | 0.99 | 2,694 |
| 303 | 7 | Blood | MNG-19 | II | 19 | 187,868 | 120,141 | 166,122 | 72.32 | 2,124.92 | 0.99 | 3,306 |
| 303 | 7 | Spleen | MNG-19 | II | 19 | 187,868 | 155,498 | 207,421 | 74.97 | 2,511.41 | 0.99 | 3,020 |
| 304 | 7 | Blood | Arm07 | II | 25 | 190,584 | 64,340 | 308,436 | 20.86 | 440.05 | 0.96 | 1,242 |
| 305 | 7 | Spleen | Arm07 | II | 21 | 190,584 | 136,574 | 173,381 | 78.77 | 2,193.54 | 0.99 | 3,035 |
| 306 | 6 | Blood | MNG-19 | II | 22 | 187,868 | 423,361 | 575,567 | 73.56 | 2,313.04 | 0.99 | 1,019 |
| 307 | 3 | Blood | MNG-19 | II | 22 | 187,868 | 264,992 | 390,731 | 67.82 | 1,465.03 | 0.99 | 1,029 |
| 308 | 7 | Blood | Arm07 | II | 21 | 190,584 | 310,219 | 426,981 | 72.65 | 1,842.86 | 0.99 | 1,117 |
| 309 | 7 | Blood | Arm07 | II | 19 | 190,584 | 96,240 | 123,894 | 77.68 | 1,726.88 | 0.99 | 3,378 |
| 310 | 7 | Blood | MNG-19 | II | 19 | 187,868 | 66,069 | 88,936 | 74.29 | 1,079.36 | 0.99 | 3,047 |
| 311 | 7 | Blood | Arm07 | II | 19 | 190,584 | 98,755 | 129,042 | 76.53 | 1,676.69 | 0.99 | 3,187 |
| 312 | 7 | Blood | Arm07 | II | 20 | 190,584 | 131,299 | 173,421 | 75.71 | 2,299.50 | 0.99 | 3,297 |
| 314 | 7 | Blood | MNG-19 | II | 22 | 187,868 | 307,810 | 421,638 | 73 | 1,714.75 | 0.99 | 1,037 |
| 511 | 7 | Blood | Arm07 | II | 20 | 190,584 | 121,736 | 152,400 | 79.88 | 1,912.97 | 0.99 | 2,970 |
| 635 | 7 | Blood | Arm07 | II | 14 | 190,584 | 165,992 | 215,116 | 77.16 | 2,654.20 | 0.99 | 3,022 |

DPC: Days post challenge; Arm07-Cos-1: ASFV Armenia 2007 passaged on Cos-1 cells; P3: Passage 3; Arm07: ASFV Armenia 2007; MNG-19: ASFV Mongolia 2019; E70: ASFV E70

**Figure 3:**
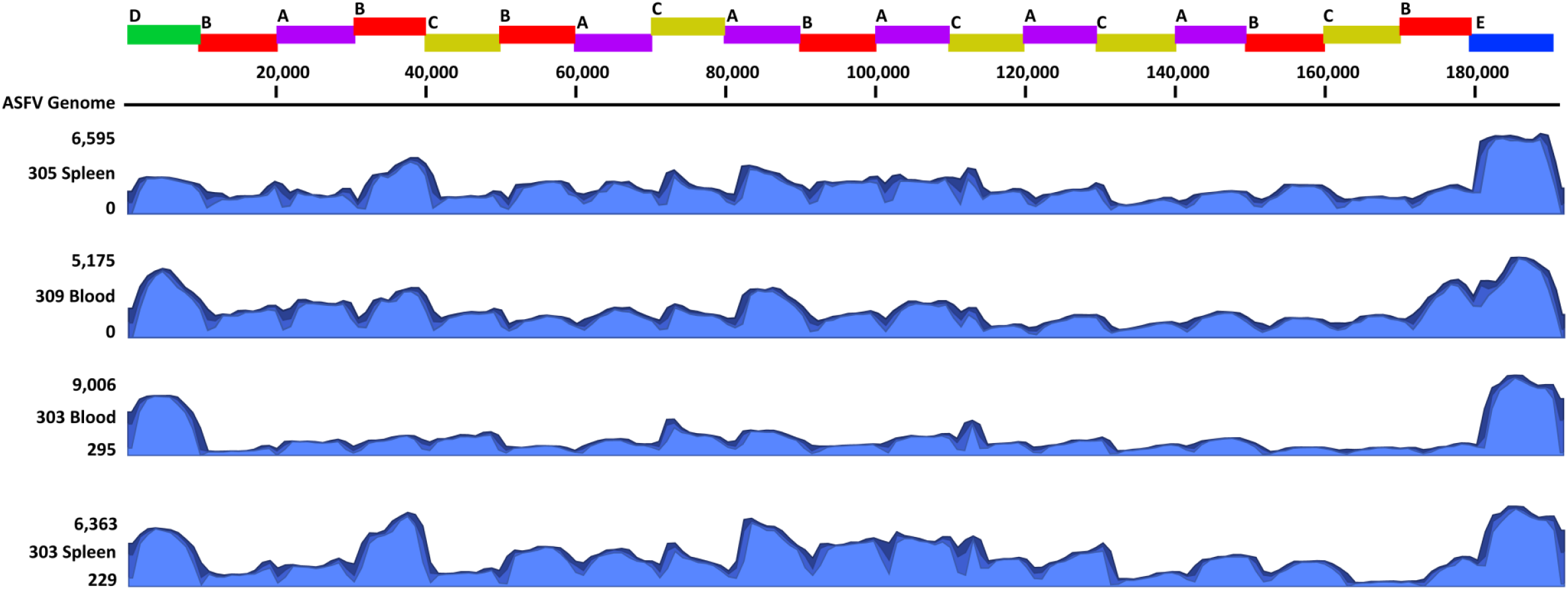
Coverage graphs of reads mapped to the ASFV genome. The primer schematic is displayed on top indicating the 19 amplicons and the associated pools such as A-E. The graphs shown are a subset of samples tested using the ASFV AmpliSeq protocol. These samples include a blood and spleen sample from a pig experimentally infected with ASFV Armenia/2007 (#305 and #309) or ASFV Mongolia/2019 (#303).

### 3.2 CSFV AmpliSeq

Two primer sets were designed to amplify approximately 6 kB fragments of all known CSFV genotypes. The primers were tested against 5 CSFV strains isolated from cell culture which included genotype 1 (Brescia, Bavaro, Guatemala), genotype 2 (Germany 99), and genotype 3 (Congenital Tremor) viruses for optimizing thermal cycling conditions and suitability to amplify viruses representing the three known genotypes. The two primer sets amplified the correct size amplicons for all test strains which were confirmed by gel electrophoresis. Smaller amplicons were observed after RT-PCR for fragment 2, likely due to partial or non-specific amplification; these smaller amplicons were excluded during the sequencing run using appropriate built-in bioinformatics filters. The efficiency of amplification for genotype 3 viruses was reduced, as observed by reduced concentrations of fragment 2 following RT-PCR. Following sequencing of the amplicons derived using the two primer sets, the average coverage ranged from 10,770X -20,356X (Figure 4 and Table 4). The large spike in coverage observed in the coverage graphs is a result of the overlapping segment between the two primer pairs. We observed a significant reduction in coverage for fragment 2 of the Congenital Tremor strain of CSFV with the depth of coverage reduced to 500-1,000 (Figure 4 and Table 4), most likely related to primer mismatch-es and/or higher sample Ct value. The CSFV AmpliSeq was also tested on 6 additional cell-derived CSFV stocks: Alfort, PAV250, Paderborn, Parma98, Germany-95, and Kanagawa (Table 4). For the 11 strains tested there was an average coverage of 12,278X with an average read length of 1.6 kB covering 97%-99% of the genome (Table 4). Similar to the Congenital Tremor strain, the other genotype 3 strain Kanagawa also had reduced amplification of fragment 2 with a reduction in coverage depth to 2,000. The reduced depth of coverage of fragment 2 for both genotype 3 strains is most likely related to primer mis-matches and/or higher Ct values. Despite the reduction in depth of coverage for fragment 2, the depth is sufficient for obtaining the near full-length genomes of genotype 3 viruses.

**Table 4.**
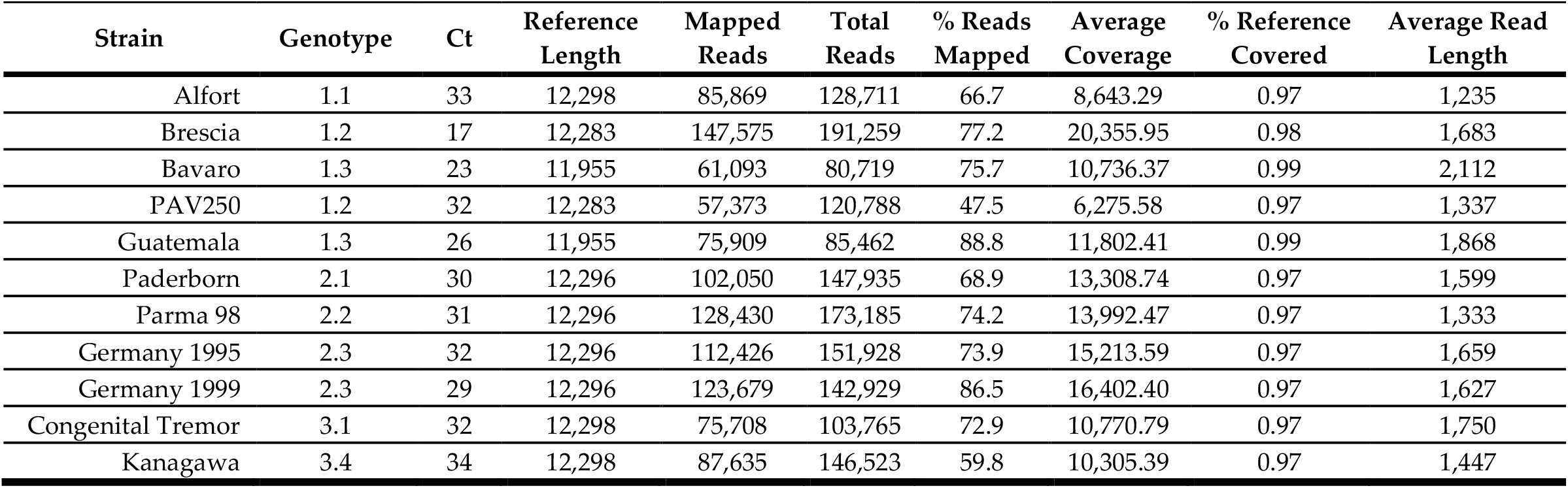
Sequencing data for 11 CSFV strains tested with CSFV AmpliSeq.

**Figure 4.**
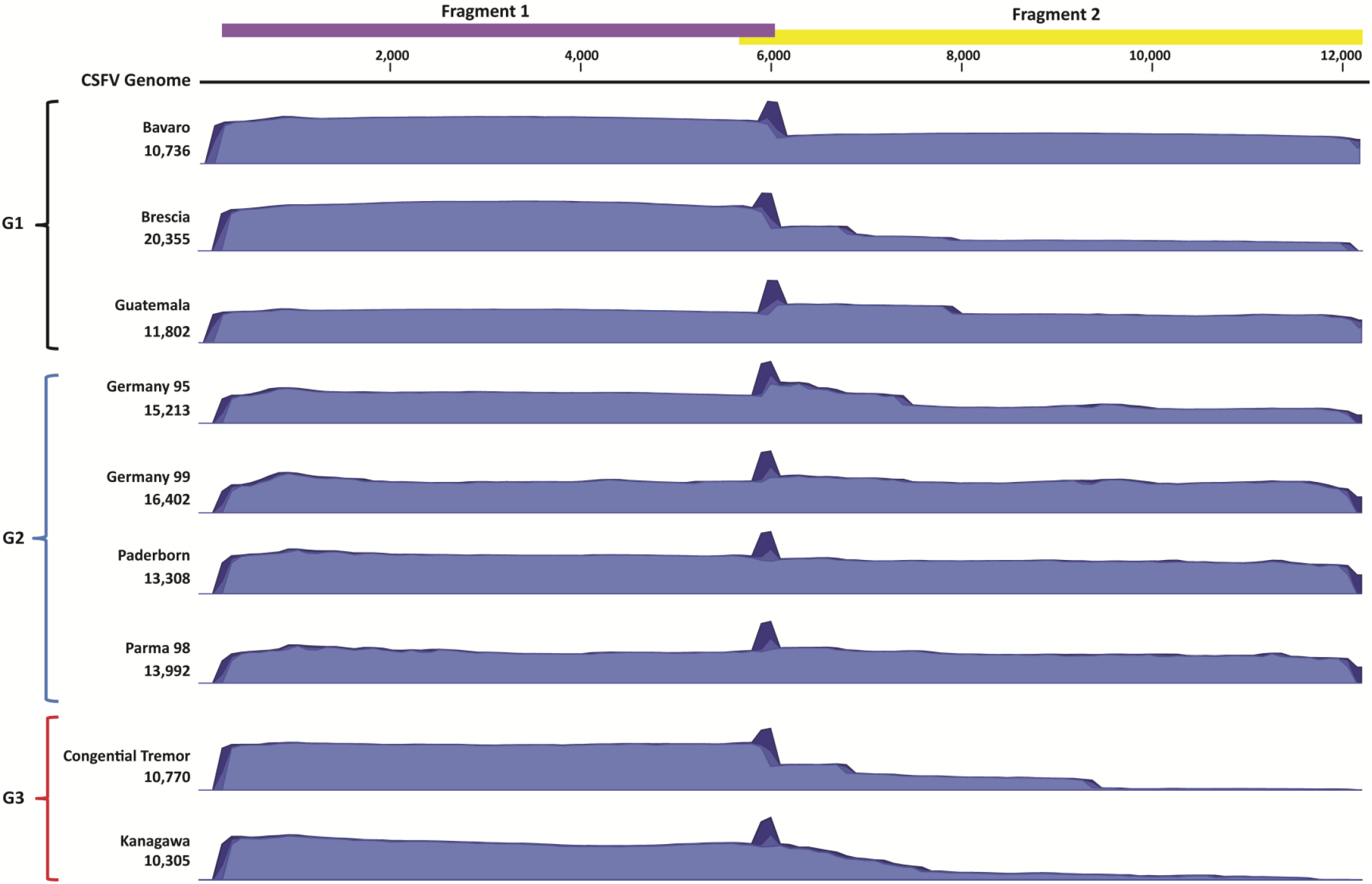
Coverage graphs for CSFV viral stocks sequenced with CSFV AmpliSeq. The primer schematic is displayed on top indicating the 2 amplicons. The graphs shown are a subset of viral stocks tested, which represent the 3 known genotypes of CSFV. Genotype 1 (G1): Bavaro, Brescia, Guatemala. Genotype 2 (G2): Germany 95, Germany 99, Paderborn, and Parma 98. Genotype 3 (G3): Congenital Tremor and Kanagawa.

To evaluate the use of this protocol for field application, we tested the protocol on clinical samples collected from pigs experimentally infected with two different genotype 1 CSFV strains, Bavaro and Brescia (see Table 5). The Ct values for these samples ranged from 16 to 29. The clinical samples included serum collected on 5 and 7 DPC, tonsils collected at 11 DPC and one mandibular lymph node collected on 5 DPC. The average coverage for the clinical samples was 15,496X with an average read length of 1.7 kB covering 98%-99% of the genome. The coverage of the clinical samples was similar to the cell culture derived virus stocks. The CSFV AmpliSeq protocol can be used to obtain near full-length CSFV genomes from clinical samples and cell-derived virus stocks representing the three currently known CSFV genotypes. While a reduction of coverage of fragment 2 was observed for genotype 3 viruses and samples with Ct values greater than 30, the depth of coverage is sufficient for phylogenetic characterization.

**Table 5.**
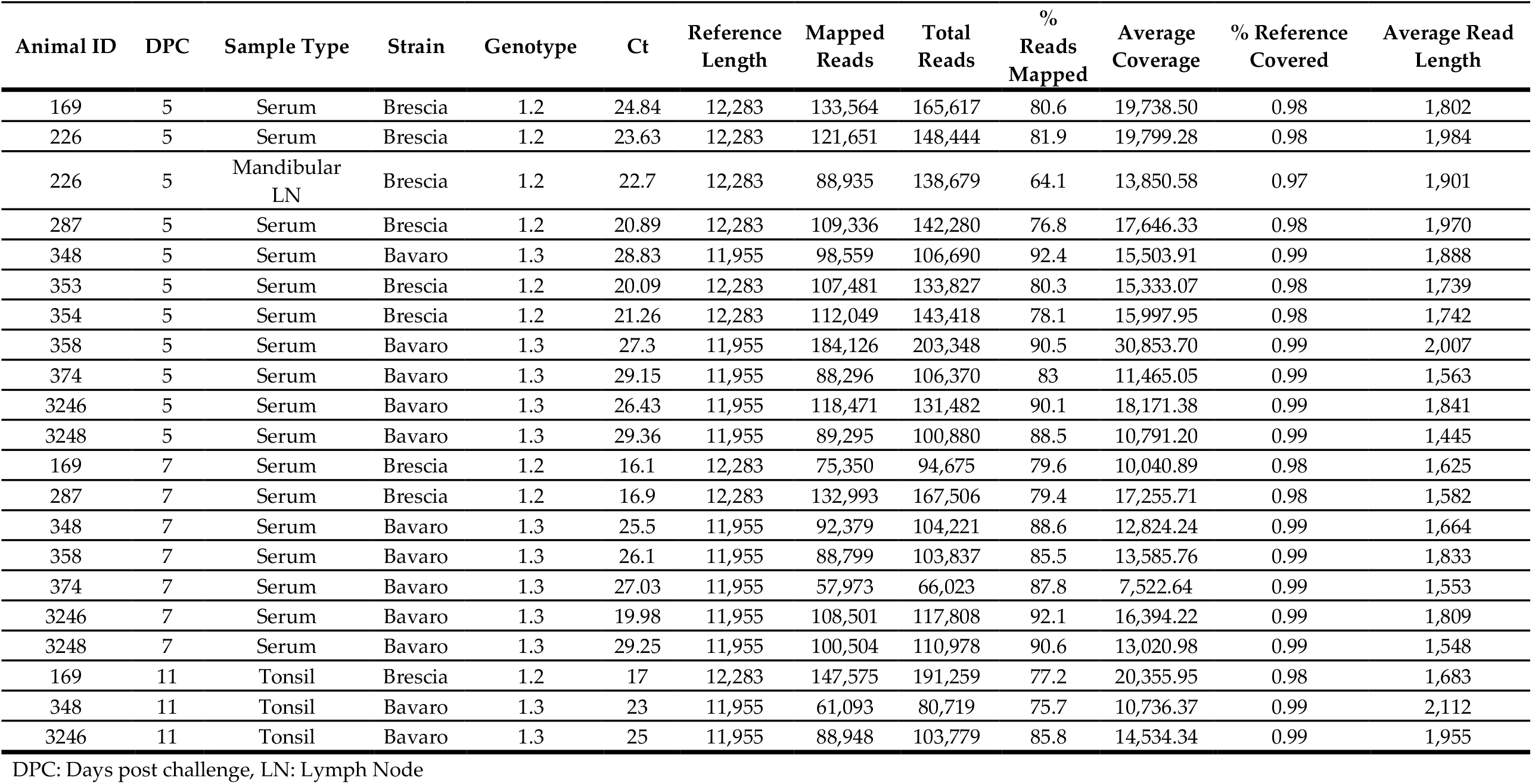
Sequencing data for clinical CSFV samples tested with CSFV AmpliSeq.

| Animal ID | DPC | Sample Type | Strain | Genotype | Ct | Reference Length | Mapped Reads | Total Reads | % Reads Mapped | Average Coverage | % Reference Covered | Average Read Length |
| --- | --- | --- | --- | --- | --- | --- | --- | --- | --- | --- | --- | --- |
| 169 | 5 | Serum | Brescia | 1.2 | 24.84 | 12,283 | 133,564 | 165,617 | 80.6 | 19,738.50 | 0.98 | 1,802 |
| 226 | 5 | Serum | Brescia | 1.2 | 23.63 | 12,283 | 121,651 | 148,444 | 81.9 | 19,799.28 | 0.98 | 1,984 |
| 226 | 5 | Mandibular LN | Brescia | 1.2 | 22.7 | 12,283 | 88,935 | 138,679 | 64.1 | 13,850.58 | 0.97 | 1,901 |
| 287 | 5 | Serum | Brescia | 1.2 | 20.89 | 12,283 | 109,336 | 142,280 | 76.8 | 17,646.33 | 0.98 | 1,970 |
| 348 | 5 | Serum | Bavaro | 1.3 | 28.83 | 11,955 | 98,559 | 106,690 | 92.4 | 15,503.91 | 0.99 | 1,888 |
| 353 | 5 | Serum | Brescia | 1.2 | 20.09 | 12,283 | 107,481 | 133,827 | 80.3 | 15,333.07 | 0.98 | 1,739 |
| 354 | 5 | Serum | Brescia | 1.2 | 21.26 | 12,283 | 112,049 | 143,418 | 78.1 | 15,997.95 | 0.98 | 1,742 |
| 358 | 5 | Serum | Bavaro | 1.3 | 27.3 | 11,955 | 184,126 | 203,348 | 90.5 | 30,853.70 | 0.99 | 2,007 |
| 374 | 5 | Serum | Bavaro | 1.3 | 29.15 | 11,955 | 88,296 | 106,370 | 83 | 11,465.05 | 0.99 | 1,563 |
| 3246 | 5 | Serum | Bavaro | 1.3 | 26.43 | 11,955 | 118,471 | 131,482 | 90.1 | 18,171.38 | 0.99 | 1,841 |
| 3248 | 5 | Serum | Bavaro | 1.3 | 29.36 | 11,955 | 89,295 | 100,880 | 88.5 | 10,791.20 | 0.99 | 1,445 |
| 169 | 7 | Serum | Brescia | 1.2 | 16.1 | 12,283 | 75,350 | 94,675 | 79.6 | 10,040.89 | 0.98 | 1,625 |
| 287 | 7 | Serum | Brescia | 1.2 | 16.9 | 12,283 | 132,993 | 167,506 | 79.4 | 17,255.71 | 0.98 | 1,582 |
| 348 | 7 | Serum | Bavaro | 1.3 | 25.5 | 11,955 | 92,379 | 104,221 | 88.6 | 12,824.24 | 0.99 | 1,664 |
| 358 | 7 | Serum | Bavaro | 1.3 | 26.1 | 11,955 | 88,799 | 103,837 | 85.5 | 13,585.76 | 0.99 | 1,833 |
| 374 | 7 | Serum | Bavaro | 1.3 | 27.03 | 11,955 | 57,973 | 66,023 | 87.8 | 7,522.64 | 0.99 | 1,553 |
| 3246 | 7 | Serum | Bavaro | 1.3 | 19.98 | 11,955 | 108,501 | 117,808 | 92.1 | 16,394.22 | 0.99 | 1,809 |
| 3248 | 7 | Serum | Bavaro | 1.3 | 29.25 | 11,955 | 100,504 | 110,978 | 90.6 | 13,020.98 | 0.99 | 1,548 |
| 169 | 11 | Tonsil | Brescia | 1.2 | 17 | 12,283 | 147,575 | 191,259 | 77.2 | 20,355.95 | 0.98 | 1,683 |
| 348 | 11 | Tonsil | Bavaro | 1.3 | 23 | 11,955 | 61,093 | 80,719 | 75.7 | 10,736.37 | 0.99 | 2,112 |
| 3246 | 11 | Tonsil | Bavaro | 1.3 | 25 | 11,955 | 88,948 | 103,779 | 85.8 | 14,534.34 | 0.99 | 1,955 |

DPC: Days post challenge, LN: Lymph Node

## 4. Discussion

Historically, ASFV and CSFV were genetically characterized and phylogenetically grouped based on partial genetic sequences due to limited sequencing capabilities. For ASFV, the community relied heavily on the partial sequencing of the major capsid protein (p72) for genotyping [16,17]. Sequencing of p54, variable region of B602L, and CD2v and C-type lectin [18,28] combined with p72 allows for further discrimination of ASFV strains. These methods provide vital information for identification and outbreak investigations but only provide limited genetic information on this large viral genome. Recent advances in sequencing technologies allowed researchers to obtain full genome sequences of ASFV through direct sequencing approaches [29]. This is a resource expensive method resulting in low coverage of the viral genome due to the overwhelming presence and sequencing of the host genome. Due to the large genome of ASFV, amplicon-based sequencing has been difficult due to the need for numerous PCR reactions. The protocol developed in this study uses an amplicon-based targeted sequencing method. We reduced the number of reactions needed to amplify the ASFV genome by generating long amplicons of approximately 10 kB; these reactions were further reduced by generating primer pools allowing for the amplification of multiple 10 kB amplicons in one reaction. The resulting protocol allows for whole genome amplification of ASFV in 5 reactions in 4 to 4.5 hours. Amplicon pools were then sequenced with the Oxford Nanopore MinION platform. With this protocol, we obtained full-length genotype II ASFV genomes in 36-48 hours with an average coverage of 1,000X at a cost of approximately $70 per sample. Along with the increased depth of coverage, this protocol provides an average error rate of 1.5% compared to the 6.5% error rate with a metagenomic approach for mapping reads to a reference genome. The protocol was also tested on a genotype I ASFV and resulted in sequences with approximately 85% of the genome covered. We concluded that the ASF AmpliSeq protocol will allow re-searchers to assemble near full-length genome sequences of this large DNA virus to study viral evolution and perform detailed phylogenetic inference.

Genetic characterization and phylogenetic analysis of CSFV historically relied on partial sequencing of the E2 gene, the 5’-NTR, or the NS5 gene [15,20]. However, with advances in sequencing technology and continuing reduction in cost for sequencing, the WOAH recommends full length sequencing of the E2 gene or whole genome sequencing. Protocols are available for amplification of the full-length E2 gene and whole CSFV genome that can be applied for sequencing [30,31]. The protocol described by Leifer et al. for amplification of the whole CSFV genome requires multiple RT-PCR reactions due to the average amplicon size being ∼1 kB. The protocol described in this study generates two ∼6 kB amplicons which serve to amplify the whole CSFV genome in two RT-PCR reactions. The pooled amplicons were sequenced on the Oxford Nanopore MinION platform. With this protocol, we obtained the near full-length genomes of viruses representing three currently known CSFV genotypes isolated from cell culture with the average coverage greater than 10,000X in 36-48 hours at a cost of approximately $30 per sample. The protocol was tested on clinical samples from pigs experimentally infected with two different genotype 1 viruses. The CSFV AmpliSeq protocol will reduce the cost and amount of sample required to obtain the whole genome sequences of CSFV, allowing for more in depth understanding of the genetic evolution of CSFV while providing the necessary genetic information re-quired for phylogenetic characterization of CSFV.

## 5. Conclusions

The targeted whole genome sequencing protocols for ASFV and CSFV described in this study will provide vital tools for diagnostics and research of these two high consequence swine pathogens that have far reaching impacts on both the global swine industry and food security. These protocols will enable researchers to rapidly obtain the near full-length genomes of ASFV and CSFV, providing more complete genetic information for phylogenetic analysis and viral evolution studies of these pathogens. Further development is needed for the ASFV AmpliSeq protocol to obtain the complete genomes of all 24 known ASFV genotypes; however, the present protocol can detect ASFV genotypes I and II that have emerged and continue to spread outside of Africa. While this study focuses on utilizing the MinION sequencing platform, these protocols would also be compatible with other sequencing platforms such as PacBio and Illumina. The reduced cost and timeframe for these protocols will be important tools to assist in early pathogen detection and genetic characterization of these high consequence swine viruses in outbreak and surveillance situations within the United States and globally.

## Author Contributions

Conceptualization of project, project administration, funding acquisition, coordination of field sample transfer, writing, review editing, J.A.R, D.G.D, R.M.P, I.M; preparation of original draft, data curation, assay development, assay optimization, sequencing, bioinformatics, C.D.M; assay development, assay optimization, review, editing, T.K; assay development, assay optimization, sequencing, review, editing, P.A; assay development, assay optimization, E.M; qPCR, data acquisition, oversite and analysis of molecular work (DNA extraction and PCR), review, and editing J.D.T; writing, review, editing, N.N.G; review and editing, L.C.C; bioinformatics, writing, editing, and review, J.S.N. All authors have read and agreed to the published version of the manuscript.

## Funding

This research was funded by United States Department of Agriculture Animal and Plant Health Inspection Services (USDA-APHIS) under agreement number AP20VSD&B000C020, the National Bio and AgroDefense Facility (NBAF) Transition Fund from the State of Kansas, the AMP and MCB Cores of the Center on Emerging and Zoonotic Infectious Diseases (CEZID) of the National Institutes of General Medical Sciences under award number P20GM130448, and the United States Department of Agriculture NBAF Scientist Training Program (USDA-NSTP) fellowship.

## Institutional Review Board Statement

The study was conducted in accordance with the Declaration of Helsinki and approved by the Institutional Biosafety Committee of Kansas State University (Protocol #s: 1651, 1668 and 1787). The animal study protocol was approved by the Institutional Animal Care and Use Committee of Kansas State University (Protocol #s: 4265 and 4363).

## Informed Consent Statement

Not Applicable.

## Data Availability Statement

Not applicable all data is included within the manuscript.

## Acknowledgments

We would like to thank the staff of the Kansas State University (KSU) Biosecurity Research Institute, the Comparative Medicine Group staff, and technical support from Sarah Lyoo, and Yonghai Lee. The authors would also like to thank Dr. Chung at Plum Island Animal Disease Center for assisting with the transfer of original samples to Kansas State University’s Biosecurity Research Institute.

## Conflicts of Interest

The J.A.R. laboratory received support from Tonix Pharmaceuticals, Genus plc, Xing Technologies, and Zoetis, outside of the reported work. J.A.R. is inventor on patents and patent applications on the use of antivirals and vaccines for the treatment and prevention of virus infections, owned by Kansas State University. The other authors declare no competing interests.

